# Genomic analysis of diet composition finds novel loci and associations with health and lifestyle

**DOI:** 10.1101/383406

**Authors:** S Fleur W Meddens, Ronald de Vlaming, Peter Bowers, Casper AP Burik, Richard Karlsson Linnér, Chanwook Lee, Aysu Okbay, Patrick Turley, Cornelius A Rietveld, Mark Alan Fontana, Mohsen Ghanbari, Fumiaki Imamura, George McMahon, Peter J van der Most, Voortman Trudy, Kaitlin H Wade, Emma L Anderson, Kim VE Braun, Pauline M Emmett, Tonũ Esko, Juan R Gonzalez, Jessica C Kiefte-de Jong, Jian’a Luan, Claudia Langenberg, Taulant Muka, Susan Ring, Fernando Rivadeneira, Josje D Schoufour, Harold Snieder, Frank JA van Rooij, Bruce HR Wolffenbuttel, 23andMe Research Team, EPIC-InterAct Consortium, Lifelines Cohort Study, George Davey Smith, Oscar H Franco, Nita G Forouhi, M Arfan Ikram, Andre G Uitterlinden, Jana V van Vliet-Ostaptchouk, Nick J Wareham, David Cesarini, K Paige Harden, James J Lee, Daniel J Benjamin, Carson C Chow, Philipp D Koellinger

**Author notes:** Correspondence and requests for materials should be addressed to SFWM, PDK, CCC, or DJB.

## Abstract

We conducted genome-wide association study (GWAS) meta-analyses of relative caloric intake from fat, protein, carbohydrates and sugar in over 235,000 individuals. We identified 21 approximately independent lead SNPs. Relative protein intake exhibits the strongest relationships with poor health, including positive genetic associations with obesity, type 2 diabetes, and heart disease (*r_g_* ≈ 0.15 – 0.5). Relative carbohydrate and sugar intake have negative genetic correlations with waist circumference, waist-hip ratio, and neighborhood poverty (|*r_g_*| ≈ 0.1 – 0.3). Overall, our results show that the relative intake of each macronutrient has a distinct genetic architecture and pattern of genetic correlations suggestive of health implications beyond caloric content.

## MAIN TEXT

Understanding the effects of nutrition on health is a priority given the ongoing worldwide obesity epidemic^1-5^. The health impacts of many aspects of dietary intake have been studied, but the effects of macronutrient composition (i.e., relative intake from fat, protein, and carbohydrate) have been especially controversial. There is still no consensus on whether macronutrients exert specific health effects beyond their caloric value^6–8^. Despite a lack of robust empirical evidence from randomized trials on the long-term effects of macronutrient restriction on body weight and health^2,6,9,10^, dietary recommendations have shifted from low-fat to low-sugar and, more recently, lower animal-protein diets^11–17^. Observational studies have found inconsistent phenotypic correlations between macronutrient proportions, body mass index (BMI) and related health outcomes (e.g., ^18–20^), and the mechanisms underlying these relationships are not well understood.

Insights from genetics may help to elucidate the connections between nutrition and health outcomes. Twin studies suggest that diet composition is moderately heritable, with *h*^2^ estimates ranging from 27% to 70% for the different macronutrients’ contributions to total energy intake^21–23^. Previous GWAS on relative caloric intake from protein, fat, and carbohydrates (up to *N* = 91,114) have identified three genome-wide significant SNPs in or near *RARB*, *FTO* and *FGF21*, each of which captures only a miniscule part of trait heritability (R^2^ < 0.06%)^24–26^. These results suggest that diet composition is a genetically complex phenotype and that most associated genetic variants have not yet been identified. Furthermore, no large-scale genome-wide association study (GWAS) has investigated relative sugar intake.

Here we report GWAS results for diet composition, and we use the results to conduct bioinformatics analyses and to calculate genetic correlations with a range of other phenotypes. For the GWAS, we expand the samples used in earlier work from *N* = 91,114^24–26^ to 268,922 for relative intake of PROTEIN, CARBOHYDRATE, and FAT. Furthermore, we report GWAS results for SUGAR (*N* = 235,391), which is a subcomponent of CARBOHYDRATE and captures relative intake of both naturally-occurring and added sugars.

## Results

### Phenotype definition

All cohorts used self-report questionnaires containing ≥70 food items, with average estimated intakes showing strong similarity across cohorts (**Supplementary Table 1.2**). Using these self-reports, we calculated the relative contributions of FAT, PROTEIN, CAARBOHYDRATE and SUGAR to total caloric intake (we do not study total caloric intake because it is mainly determined by body size and physical activity^27^ and because systematic underreporting of total food intake is correlated with BMI^28^). Since macronutrient intake may not scale linearly with total caloric intake, we developed and applied a method that adjusts for the observed non-linear relationships (**Supplementary Information 2.6, Extended Data Figure 1**). Consistent with the satiating properties of protein^29^, we find that at higher levels of total caloric intake, relative protein intake declines, while relative fat intake increases, and relative sugar and carbohydrate intake remain roughly constant (**Supplementary Table 2.2**).

### Main results

We began by assessing the SNP-based heritability of our phenotypes. We calculated GREML^30^ estimates using a random *N* = 30,000 subsample of conventionally unrelated UK Biobank (UKB) individuals. The estimates range from 2.1% for PROTEIN to 7.9% for CARBOHYDRATE (**Extended Data Figure 2** and **Supplementary Information 7**).

GWAS were performed in individuals of European ancestry. When possible, we excluded individuals on calorie- or macronutrient-restricted diets (**Supplementary Table 1.3**). Our discovery sample was the subset of the UKB with survey data on dietary intake (*N* = 175,253). The replication phase consisted of a meta-analysis of GWAS summary statistics from 14 additional cohorts that followed our analysis plan (*N* = 60,138) and summary statistics from DietGen^25^ (for FAT, PROTEIN and CARBOHYDRATE, *N* = 33,531). DietGen^25^ assumed a linear scaling of macronutrients with total energy intake. Since the genetic correlations between DietGen and our replication cohorts is not significantly different from 1 (**Supplementary Table 6.1**), we added DietGen to our meta-analysis.

Association statistics underwent rigorous quality control (**Supplementary Information 3.3**). The discovery stage identified 21 approximately independent genome-wide-significant lead SNPs (see **Supplementary Information 3.3.5** for a description of the clumping algorithm): 4 for FAT, 5 for PROTEIN, 5 for SUGAR, and 7 for CARBOHYDRATE (**Supplementary Table 4.1**). These lead SNPs partially overlap across phenotypes and reside in 14 unique loci. In the replication stage, all 21 lead SNPs had the anticipated signs and comparable effect sizes (**Extended Data Figure 3**), and 15 reached statistical significance at *P* < 0.05 (**Supplementary Table 4.1**). This empirical replication record matches or exceeds theoretical predictions that take into account the statistical winner’s curse, sampling variation, and statistical power^31^ (**Supplementary Information 4.1**).

In order to maximize statistical power, all follow-up analyses that now follow are based on results from the combined analyses of discovery and replication samples (*N* = 235,391 to 268,922). The quantile-quantile plots of exhibit substantial inflation (λ_GC_ = 1.12 to 1.19, **Extended Data Figure 4**). The estimated intercepts from LD Score regressions^32^ (LDSC) suggest that the vast majority of this inflation is due to polygenic signal, and only a small share is attributable to population stratification (max ~6% for FAT, *n.s*. different from 0%; **Supplementary Table 3.4**). The number of approximately independent lead SNPs is 36 (pairwise *r^2^* < 0.01), including 6 for FAT, 7 for PROTEIN, 10 for SUGAR, and 13 for CARBOHYDRATE (**Table 1, Figure 1**). These 36 lead SNPs reside in 21 unique loci (**Supplementary Table 5.2**). The SNP effect sizes range from 0.015 to 0.098 phenotypic standard deviations per allele. The phenotypic variance explained per SNP, expressed in terms of coefficient of determination (*R^2^*), ranged from 0.011% to 0.054%, comparable to other genetically complex traits such as BMI and educational attainment (**Extended Data Figure 5**).

**Figure 1.**
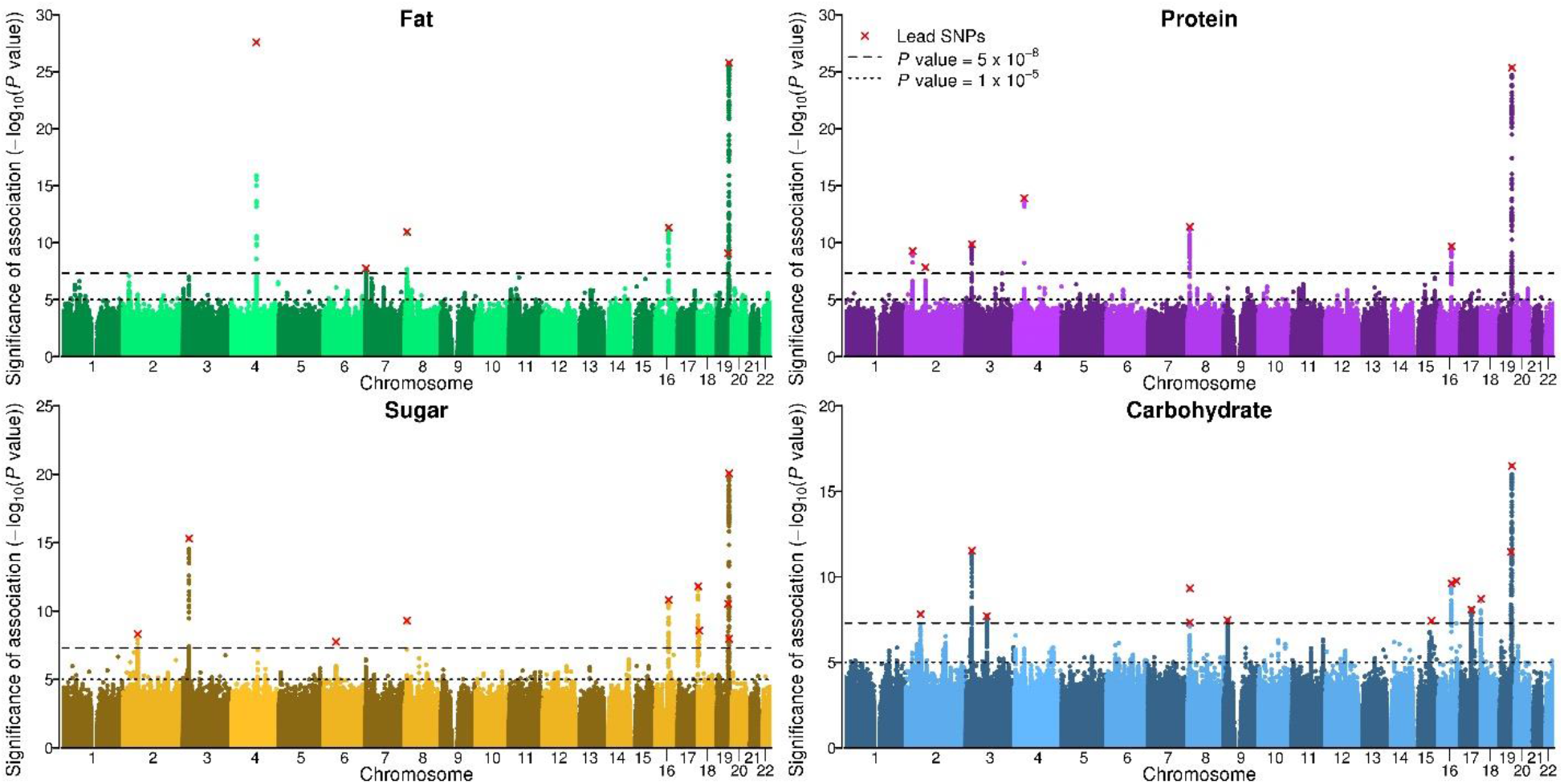
Manhattan plots. The *x*-axis is SNP chromosomal position; the *y*-axis is the SNP *P* value on a −log_10_ scale; the horizontal dashed line marks the threshold for genome-wide (*P* = 5 × 10^−8^) and suggestive (*P* = 1 × 10^−5^) significance; and each approximately independent (pairwise *r^2^* < 0.1) genome-wide significant association (“lead SNP”) is marked by a red cross.

**Table 1.**
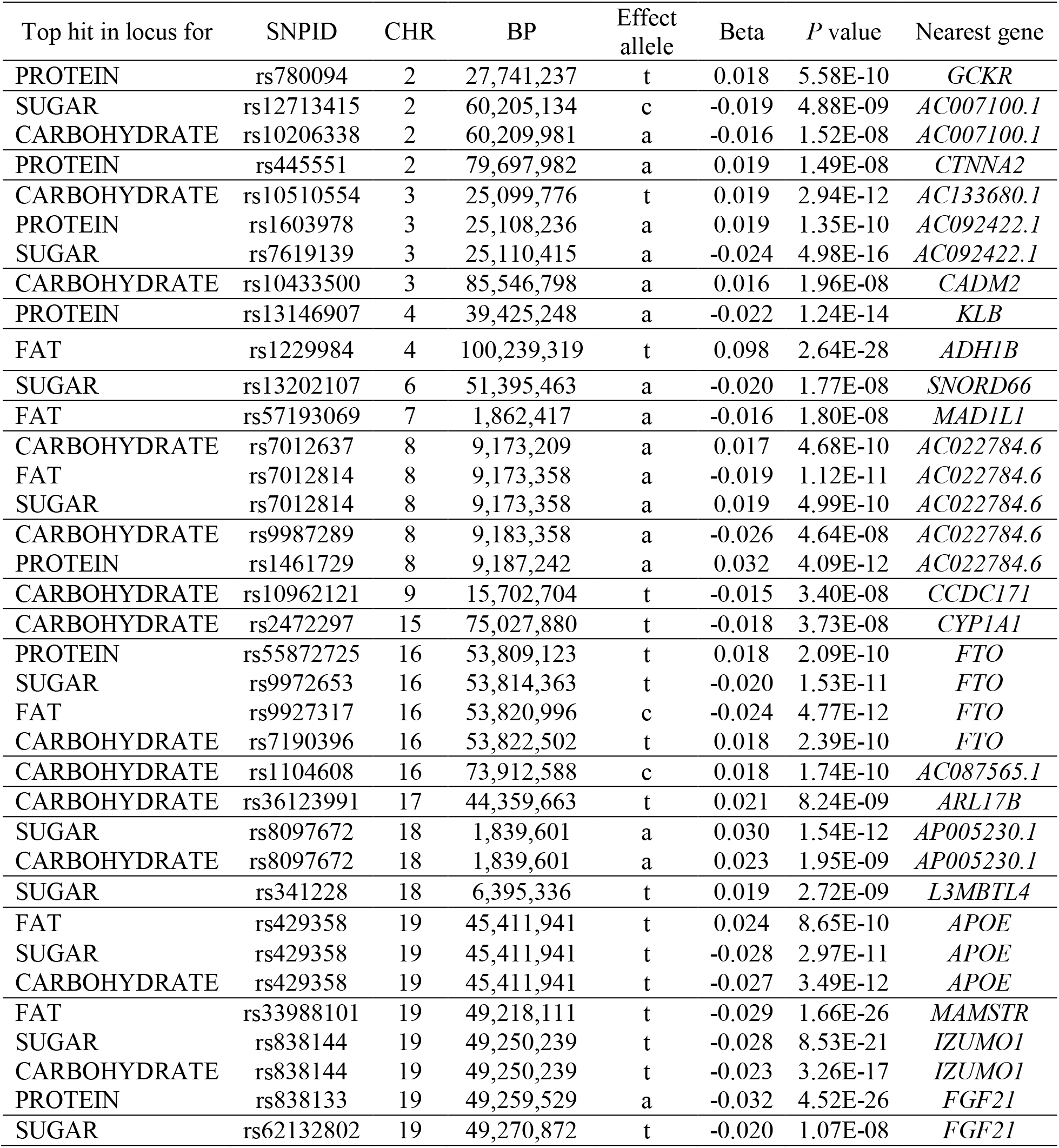
Diet composition lead SNPs. GWAS summary statistics of the 36 diet composition lead SNPs (i.e., the top hit in the locus for each phenotype). A total of 21 of these lead SNPs are approximately independent. **Supplementary Table 5.1** reports the effect alleles and summary statistics across all four phenotypes for each individual lead SNP. MAF = minor allele frequency (weighted average across cohorts). Beta = semi-standardized (i.e., increase in phenotypic standard deviations per effect allele). All *P* values are genomic-controlled (corrected for LDSC intercept). All genomic coordinates are in GRCh37.

MAGMA^33^ analyses of our GWAS summary statistics identified 81 unique genes (**Extended Data Figure 6** and **Supplementary Table 5.4**). While the majority of these genes were near our lead SNPs, MAGMA also identified 33 genomic regions harboring 44 unique genes that are physically distant (> 1 Mb) from our lead SNPs.

We constructed polygenic scores for the macronutrient intakes by applying LDpred^34^ to our GWAS summary statistics. We assessed the scores’ out-of-sample predictive accuracy in two holdout cohorts: the Health and Retirement Study (*N* = 2,344) and the Rotterdam Study (*N* = 3,585). The scores predicted the macronutrient intakes with *R*^2^ ranging between 0.08% (*P* = 0.088) and 0.71% (*P* = 9.11×10^−7^; **Supplementary Table 8.1**).

We estimated pairwise genetic correlations between the macronutrients with bivariate LDSC^35^. All are statistically distinguishable from zero at *P* < 0.05 (except FAT and PROTEIN), as well as from one and negative one (**Table 2**). These results indicate that intake of each macronutrient has a different genetic architecture, consistent with previous work from animal studies showing distinct biological mechanisms involved in macronutrient-specific appetites^36^.

**Table 2.** Genetic correlations between macronutrients. Genetic correlation analysis results obtained from bivariate LD Score regression (with block jackknife standard errors in brackets). Only HapMap3 SNPs were used in this analysis. The results show the genetic correlations among the four phenotypes calculated using the summary statistics from the combined meta-analyses. *** Denotes *P* value < 0.001 for the null hypothesis of zero genetic correlation. All estimates also differed from 1 and −1 with *P* < 0.001.

|  | FAT | PROTEIN | Sugar | CARBOHYDRATE |
| --- | --- | --- | --- | --- |
| FAT | -- | -- | -- | -- |
| PROTEIN | -0.019 (0.068) | -- | -- | -- |
| SUGAR | -0.513 (0.040)*** | -0.307 (0.057)*** | -- | -- |
| CARBOHYDRATE | -0.607 (0.032)*** | -0.226 (0.048)*** | 0.728 (0.020)*** | -- |

### Discussion of lead SNPs from combined meta-analysis

Seven of the 21 lead SNPs have not been (directly or via LD partners, *r*^2^ ≥ 0.6 and distance < 250 kb) associated with any other traits in the NHGRI-EBI GWAS Catalog^37^ (**Supplementary Table 5.5**). Each of these seven SNPs is located in or near genes that have not been studied in depth to date.

Five lead SNPs are located in (or near) genes that have well characterized biological functions in nutrient metabolism or homeostasis but have not previously been associated with food intake. First, a missense variant in *APOE* (rs429358) was associated with FAT, SUGAR, and CARBOHYDRATE, where the allele that decreases Alzheimer’s risk is associated with greater FAT intake, and vice-versa for SUGAR and CARBOHYDRATE. *APOE* is not only strongly associated with Alzheimer’s disease^38^ but is also involved in fatty acid metabolism. We explored whether this association may be driven by sample selection. Specifically, older people with dementia may be systematically missing from the UKB, and unaffected elderly people may have different eating habits than younger people. We found that the association was greatly reduced in the subsample of UKB participants aged below 60, but the 95% confidence intervals of the effect sizes still overlapped with those of the older sample (**Supplementary Table 5.3**).

Second, a well-known missense variant (rs1229984 in *ADH1B*) that limits alcohol metabolism was positively associated with FAT intake. The association was weaker in a sample of UKB alcohol abstainers (*N* = 39,679; **Supplementary Table 5.3**), suggesting that it may be partially driven by substitution of fat for alcohol.

Third, a protein lead SNP (rs13146907) was found in *KLB*, an essential cofactor to FGF21^39,40^ which influences sweet and alcohol taste preference via the liver-brain-endocrine axis^41–43^. *KLB* was only associated with PROTEIN, while variants in (or near) *FGF21* were strongly associated with all four macronutrients. With MAGMA, we also identified *MLXIPL* (only for FAT), a gene that acts as a transcription factor to FGF21^44^. This might imply that different genes involved in the same pathway are important for directing intake of different macronutrients.

Fourth, an intergenic variant (rs2472297) linked to higher caffeine consumption^45,46^ was associated with lower CARBOHYDRATE intake. There are various possible explanations, such as interrelated lifestyle choices pertaining to food and caffeinated drinks.

Fifth, an intronic variant in *GCKR* (rs780094), a carbohydrate-metabolism gene, is associated with PROTEIN. The lead SNP is in almost perfect LD with a missense variant that has been associated with lipid levels^47^.

### Bioinformatic analyses

Animal studies indicate that the brain and peripheral organs interact in directing macronutrient intake^36,48^. A question that arises is whether the “periphery”, which digests and metabolizes macronutrients, plays a larger role than the brain, for instance by determining how the brain assigns reward values to macronutrients. (For example, this is partially the case with alcohol, where mutations that limit metabolic capacity render alcohol consumption unpleasant^49,50^.) To examine to what extent genetic variation in the brain and the periphery contributes to macronutrient intake in humans, we used stratified LDSC^51,52^ to identify in which tissues diet-composition-associated SNPs are likely to be expressed (**Supplementary Information 9.1**). We performed two stratified LDSC analyses, which partitioned SNP heritability according to (i) 10 broadly-defined tissues, which were ascertained with LDSC reference data from chromatin data^53^ and (ii) 53 tissues (including 14 brain regions), as ascertained with LDSC reference data from sets of Specifically Expressed Genes in GTEx (known as LDSC-SEG)^52^. To correct for multiple testing across tissues, we applied Bonferroni adjustments for the number of tested tissues (*P*_Bonf_ = 10 · *P* and *P*_Bonf_ =53 · *P*, respectively).

We found that genetic variation related to the central nervous system plays a major role for intake of all macronutrients (*P*_Bonf_ < 0.015 for the regression coefficients; Figure 4), with the proportions of explained heritability ranging from 44% (FAT and SUGAR) to 55% (PROTEIN). Within the central nervous system, we found broad involvement of the brain, including (frontal) cortex (FAT and SUGAR), the basal ganglia (FAT), limbic system (FAT and SUGAR), cerebellum (PROTEIN), and hypothalamus and substantia nigra for FAT and PROTEIN (and SUGAR suggestively: *P*_Bonf_ = 0.06). The confidence intervals for the coefficients overlap across brain regions so we cannot draw conclusions about the specificity of brain regions for intake of particular macronutrients.

For FAT, genetic variation related to adrenals and/or pancreas tissue is estimated to explain 37% of the heritability. Because the adrenals play a role in lipid metabolism, and the pancreas is crucial for digestion, either tissue may plausibly affect fat intake. We caution, however, that in the LDSC-SEG analyses of 53 tissues, all non-brain regions had *P* values above 0.05 even before Bonferroni adjustment (Figure 5).

To gain insight into the putative functions of the top associated loci, we queried the 81 genes identified by the MAGMA analyses in Gene Network^54^, which predicts Reactome^55^ functions for genes (**Supplementary Information 9.3**). In addition to neural functioning (e.g., axon guidance), we found that the MAGMA genes were predicted to be involved in growth factor signaling and the immune system (**Supplementary Table 9.6**). These results may imply a more pronounced role for peripheral gene functions than our stratified LDSC results, which mainly implicated the brain.

**Figure 4.**
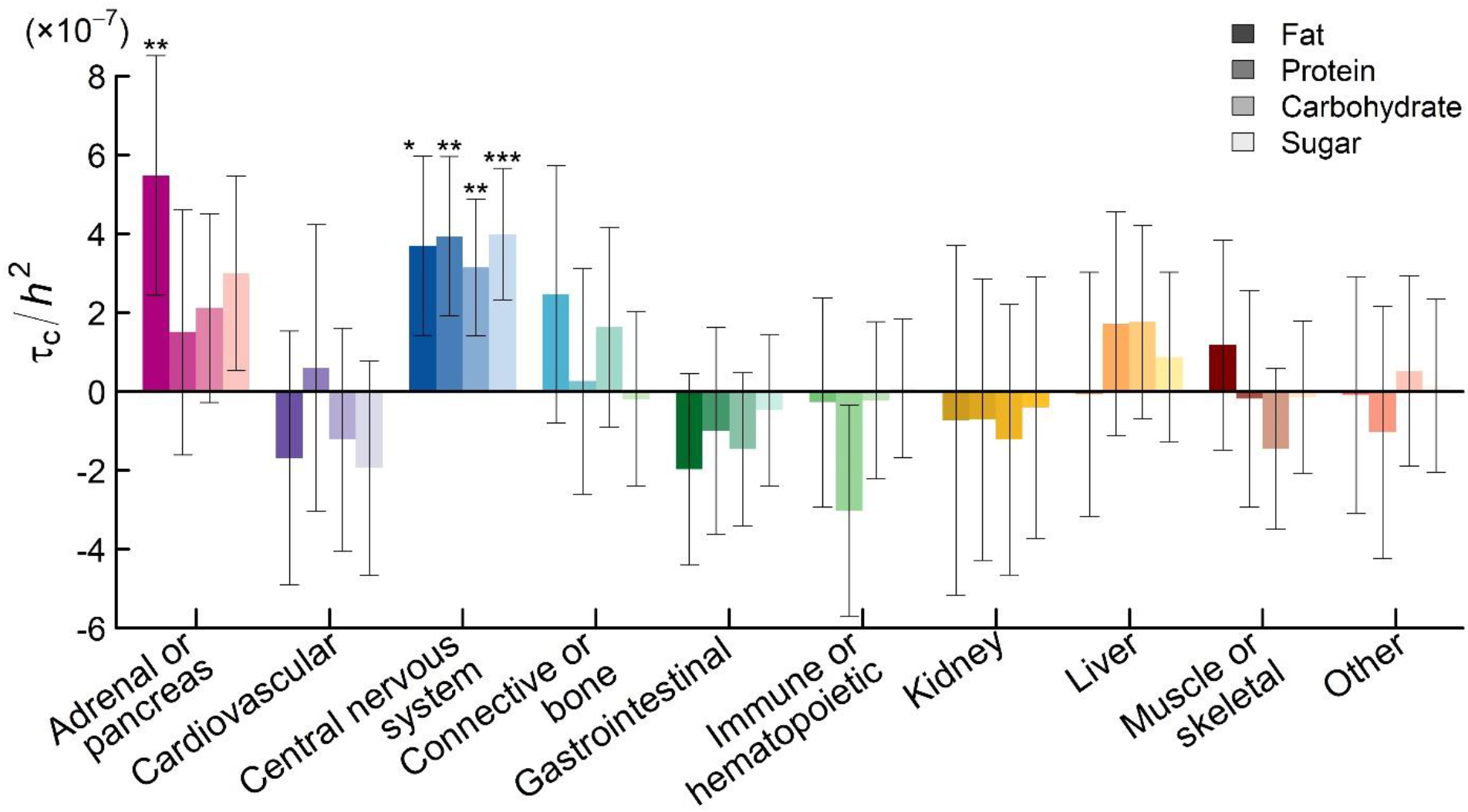
LD Score partitioning of heritability – Tissues. Functional partitioning of the heritability of diet composition phenotypes with stratified LD Score regression, where tissues were ascertained by Finucane et al. on the basis of chromatin data. The panel shows the partial regression coefficient (*τ_C_*) from the stratified regression, divided by the LD Score heritability of the diet composition phenotype (*h^2^*). Each estimate of *τ_C_* comes from a separate stratified LD Score regression, where we also controlled for the 52 functional annotation categories in the “baseline” model. Error bars represent 95% confidence intervals. The phenotypes are ordered from left to right (FAT, PROTEIN, SUGAR and CARBOHYDRATE), from darker to lighter shades. Asterisks (*) denote significant deviation from zero after Bonferroni correction for 10 tissues: * 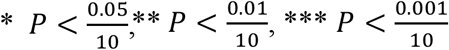.

**Figure 5.**
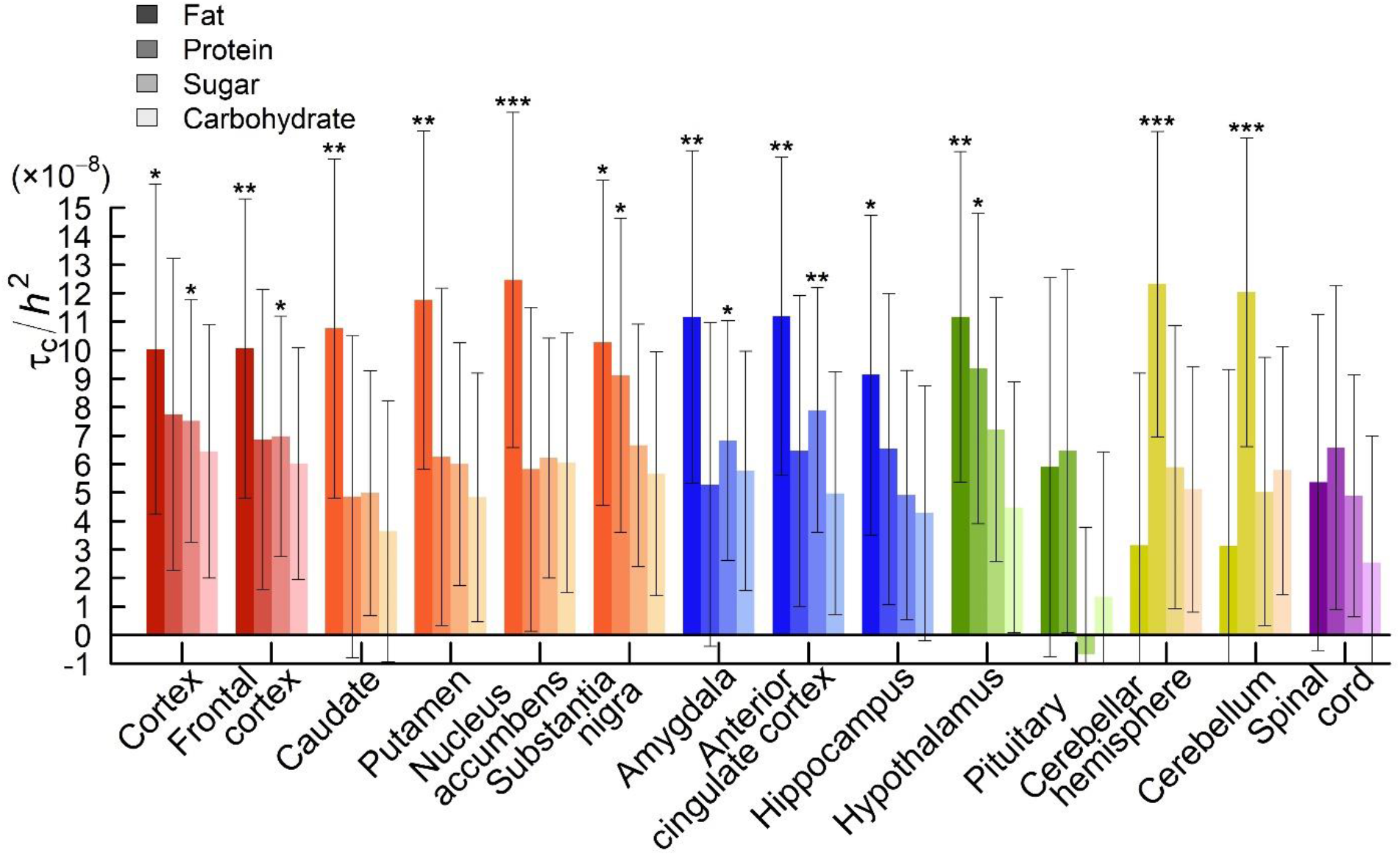
LD Score partitioning of heritability – Brain regions. Functional partitioning of the heritability of diet composition phenotypes with stratified LD Score regression, where tissues were ascertained by Finucane et al. on the basis of sets of specifically-expressed genes in GTEx data (“LDSC-SEG”). The sets of specifically-expressed genes in these analyses compared the focal tissue to other bodily tissues. The panel shows the partial regression coefficient (*τ_C_*) from the stratified regression, divided by the LD Score heritability of the diet composition phenotype (*h^2^*) to facilitate comparison between traits. Each estimate of *τ_C_* comes from a separate stratified LD Score regression, where we also controlled for the 52 functional annotation categories in the “baseline” model. Error bars represent 95% confidence intervals. Asterisks (*) denote significant deviation from zero after Bonferroni correction for 53 tissues; 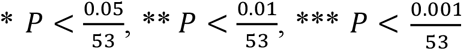. Each group of colored bars represents an anatomical region (ordered from left to right: red – cortex, orange – basal ganglia, blue – limbic system, green – hypothalamus-pituitary, yellow – cerebellum, and purple – spinal cord).

### Relationships with health, lifestyle and socioeconomic status

Using bivariate LDSC^35,56^, we estimated genetic correlations between our diet-composition phenotypes and 19 preselected relevant medical and lifestyle phenotypes for which well-powered GWAS results were available. We also included four additional phenotypes for which GWAS results became available after our study was underway, as well as Alzheimer’s disease, motivated by the association we found between *APOE* with macronutrient intake. To control for multiple testing, we again calculated Bonferroni-adjusted *P* values (*P*_Bonf_ = 24 · *P*).

PROTEIN showed the strongest genetic correlations with poor health outcomes, including obesity (*r_g_ =* 0.35), type 2 diabetes (*r_g_* = 0.45), fasting insulin (*r_g_* = 0.41), and coronary artery disease (*r_g_* = 0.16), as well as BMI (*r_g_* = 0.40) (**Figure 2, Supplementary Table 10.1**). FAT, SUGAR, and CARBOHYDRATE had negative, non-significant genetic correlations with BMI (*r_g_* between −0.06 and −0.02). For comparison, we estimated phenotypic associations between diet composition and BMI in four independent cohorts (combined *N* = 173,353) and meta-analyzed the results (Figure 3). PROTEIN (standardized 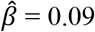) and FAT (standardized 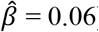) are positively associated with BMI, while SUGAR and CARBOHYDRATE are negatively associated with BMI (standardized 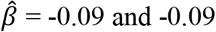 and −0.09, respectively, **Supplementary Table 10.2**). Thus, the genetic correlation between PROTEIN and BMI stands out as large relative to the phenotypic correlation.

**Figure 2.**
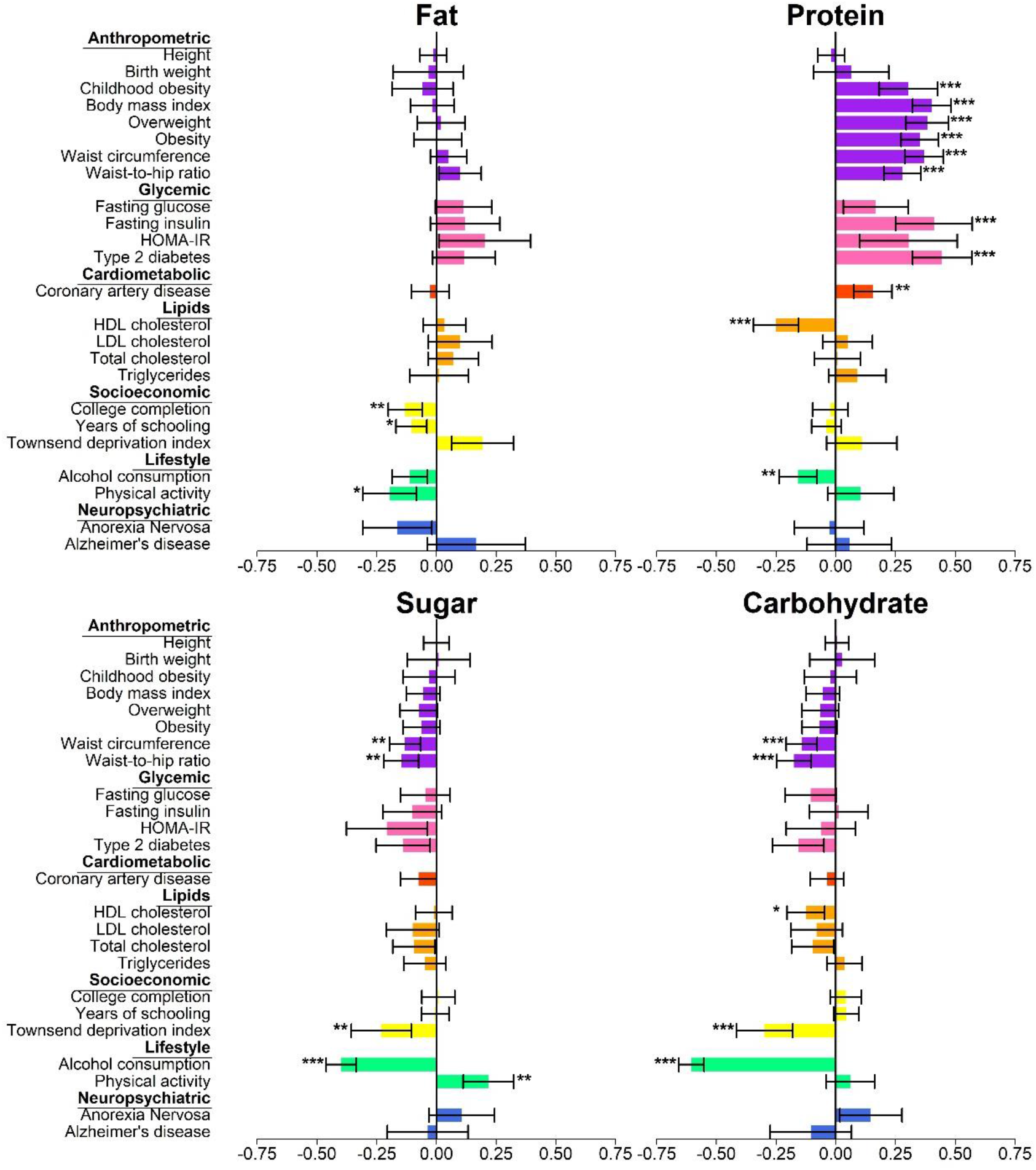
Genetic correlations. Genetic correlations were estimated with bivariate LD Score (LDSC) regression. Error bars show 95% confidence intervals, while asterisks denote Bonferroni-corrected *P* value thresholds (* *P*_Bonferroni_ < 0.05, ** < 0.01, *** < 0.001), corrected for 24 traits. The colours represent the different functional domains.

**Figure 3.**
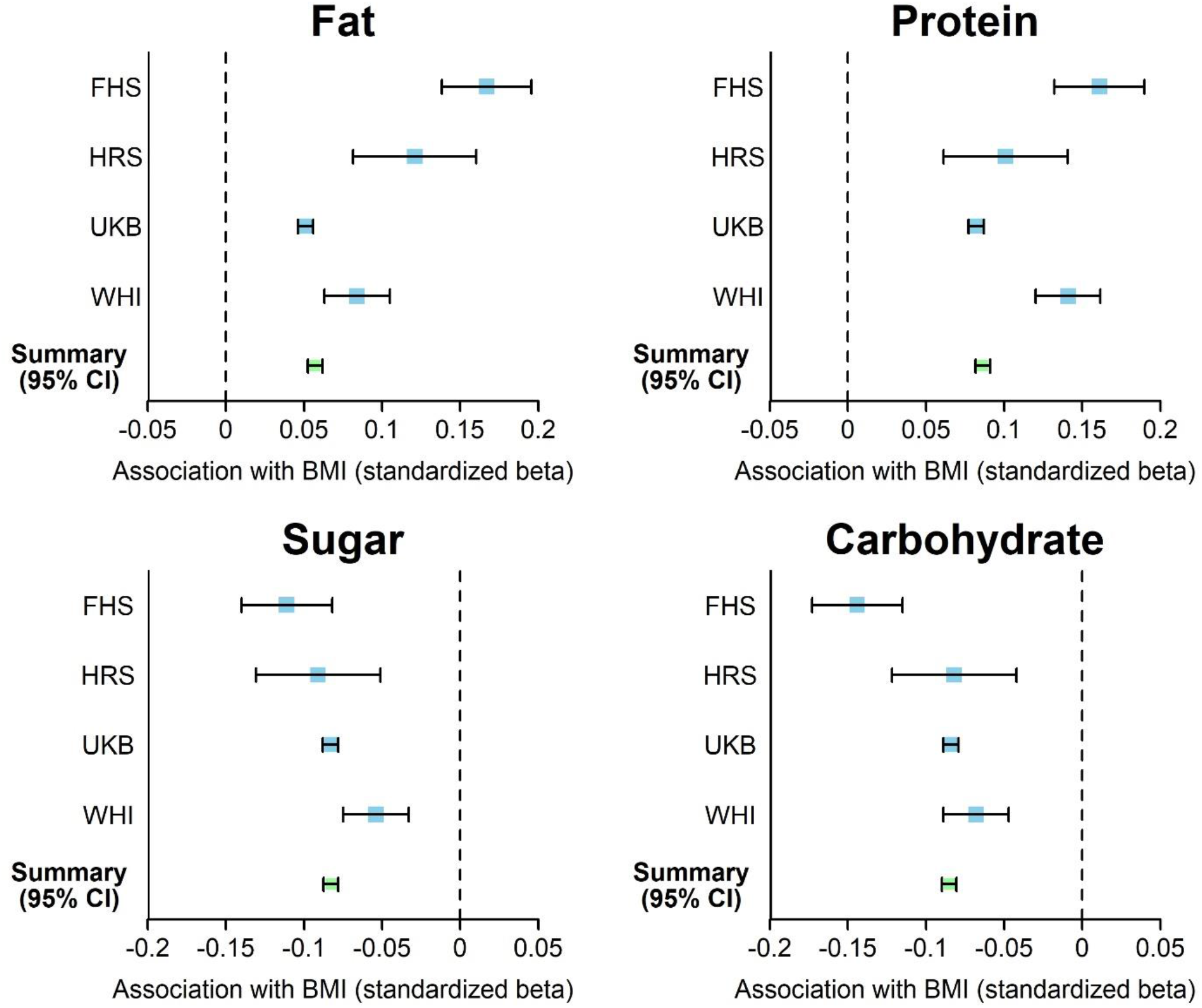
Phenotypic associations with Body Mass Index. Forest plots depicting the phenotypic associations between diet composition and Body Mass Index (BMI) in four independent cohorts, in terms of standardized betas (with errors bars indicating 95% confidence intervals). These standardized regression coefficients were obtained from a regression of BMI on the focal macronutrient and several covariates (sex, age, educational attainment, and household income). FHS = Framingham Heart Study (*N* = 4,413), HRS = Health and Retirement Study (*N* = 2,394), UKB = UK Biobank (*N* = 158,046), WHI = Women’s Health Initiative (*N* = 8,628). The summary estimate was based on a fixed-effects, inverse-variance weighted meta-analysis of all four cohorts.

Despite their relatively weak genetic correlations with BMI, SUGAR and CARBOHYDRATE have significant negative genetic correlations with waist circumference (*r_g_* = −0.13 and −0.14) and waist-hip ratio (*r_g_* = −0.15 and −0.18). All the macronutrients have negative genetic correlations with alcohol consumption (¾ between −0.61 and −0.11), as expected since alcohol is included in energy intake and our phenotype measures are shares of energy intake.

Next, we computed genetic correlations with indicators of socioeconomic status^31,57,58^, which are known to be phenotypically associated with food access, dietary choices, and health^59–63^. We found that FAT is negatively genetically correlated with educational attainment (*r_g_* = −0.13). SUGAR and CARBOHYDRATE are negatively genetically correlated with the Townsend deprivation index (¾ = −0.23 and −0.30), which is constructed from the rates of unemployment, non-ownership of cars and houses, and overcrowding of the neighborhood in which individuals live^64,58^, with higher scores indicating more severe socioeconomic deprivation. These genetic correlations are suggestive of environmental channels that affect macronutrient intake.

Finally, we estimate the genetic correlations between diet composition and physical activity. Because physical activity is known to have health benefits^65^, its genetic correlations with diet composition may provide clues about mechanisms underlying relationships between diet composition and health. In these genetic correlation analyses, we used unpublished physical activity GWAS summary statistics from a sample of research participants from 23andMe (*N* = 123,983). The physical activity phenotype is a composite measure based on self-reported activities from leisure, occupation, and commuting. We found a negative genetic correlation of physical activity with FAT (*r_g_* = −0.20) and a positive genetic correlation with SUGAR (*r_g_* = 0.22). The genetic correlations with PROTEIN and CARBOHYDRATE are positive but not statistically distinguishable from zero (0.11 and 0.06, respectively).

## Discussion

A possible role for PROTEIN in the etiology of metabolic dysfunction is implicated by the genetic correlation between PROTEIN and obesity, waist-hip ratio, fasting insulin, type 2 diabetes, HDL cholesterol, and heart disease, as well as by the BMI-increasing *FTO* allele associating with increased protein intake. This conclusion coincides with a growing (but often overlooked^66^) body of evidence that links protein intake to obesity and insulin resistance^67–75^. The positive genetic link between PROTEIN and BMI could reflect a causal effect of relative protein intake. There is some evidence from randomized trials with infants, which found a causal relationship between high-protein baby formula and infant body fat^76^. While the underlying biological mechanisms are unclear, high consumption of protein or certain types of amino acids (i.e., building blocks of protein) can induce insulin resistance^77–79^, rapamycin signaling^72^, and growth factor signaling^80^, thereby increasing metabolic dysfunction and early mortality risk.

We caution, however, that the strong and consistent links between PROTEIN and poor health outcomes might also be consistent with alternative explanations. Causation could run in the reverse direction: overweight individuals may have higher protein needs, or use high-protein diets as a weight-loss strategy. The associations might also be caused by other, unmeasured variables such as unhealthy lifestyle factors or co-consumed ingredients. However, we find that the phenotypic association between PROTEIN and BMI is robust to controls for educational attainment and household income. Furthermore, the genetic correlation between PROTEIN and physical activity is statistically indistinguishable from zero but positive. These findings weigh against socioeconomic status or physical activity being confounders of the positive genetic correlation between PROTEIN and BMI.

For SUGAR, the phenotypic and genetic correlations we found with BMI and other health outcomes are consistent with observations from systematic reviews and meta-analyses of phenotypic relationships. Together, this body of evidence suggests that dietary sugar, beyond its caloric value, does not have negative health effects^81–85^, contrary to some popular beliefs (e.g., ^17^). Another possibility is that exercise offsets negative metabolic effects of high sugar intake^86,87^. Those with a higher predisposition to be physically active may tend to consume more sugar, as sugar is a metabolically convenient source of energy during exercise^88^ and may enhance endurance^89^. If so, the positive genetic correlation between SUGAR and physical activity might partially explain the lack of genetic correlations between SUGAR and poor health.

For FAT and CARBOHYDRATE, we also found no consistent pattern of genetic and phenotypic associations with poor metabolic health. Taken together, our results complement the findings of phenotypic analyses from a large, multinational study by the EPIC-PANACEA consortium (*N* = 373,803), which found that only calories from protein are associated with prospective weight gain^18^ – a finding that was consistent across 10 countries.

While the phenotypic associations between dietary intake and health and lifestyle factors have been extensively explored in prior work, the large-scale genetic study of dietary intake is new. Overall, our results show that the relative intake of each macronutrient has a distinct genetic architecture, and the pattern of genetic correlations is suggestive of health implications beyond caloric content. Moreover, our genetic correlation and bioinformatics analyses suggest a number of novel hypotheses regarding the causes and consequences of dietary intake that can be explored in future work.

## Online Methods

Materials and methods are described in detail in the online Supplementary Information. Upon publication, GWAS summary statistics for the four macronutrients can be downloaded from the SSGAC website (https://thessgac.org/data).

## Acknowledgements

This research was carried out under the auspices of the Social Science Genetic Association Consortium (SSGAC, https://www.thessgac.org/. The research has also been conducted using the UK Biobank Resource under Application Number 11425. The study was supported by funding from the Ragnar Söderberg Foundation (E9/11 and E42/15), the Swedish Research Council (421-2013-1061), The Jan Wallander and Tom Hedelius Foundation, an ERC Consolidator Grant to Philipp Koellinger (647648 EdGe), the Pershing Square Fund of the Foundations of Human Behavior, The Open Philanthropy Project (2016-152872), and the NIA/NIH through grants P01-AG005842, P01-AG005842-20S2, P30-AG012810, and T32-AG000186-23 to NBER and R01-AG042568-02 to the University of Southern California. CCC was supported by the Intramural Research Program of the NIH/NIDDK. We thank the DietGen and CHARGE consortia for sharing diet composition GWAS summary statistics, and we thank 23andMe, Inc., for sharing physical activity GWAS summary statistics. A full list of acknowledgements is provided in **Supplementary Information 13**.

## Author Contributions

DJB, CCC, PDK, and SFWM designed and oversaw the study. RdV proposed the phenotype construction. SFWM was the lead analyst, responsible for GWAS, quality control, meta-analysis, summarizing the overlap across the results of the various GWAS, heritability analyses, genetic correlation analysis, phenotypic correlation analyses, out-of-sample prediction, and all bioinformatics analyses. RKL assisted with GWAS in UKB, and RKL and AO assisted with cohort-level quality control. CL performed the replication analyses, and was supervised by PT and DJB. CAPB performed the impG imputation of DietGen summary statistics. SFWM prepared the majority of figures with assistance from RKL; PB and CL also prepared some figures. JJL provided helpful advice and feedback on various aspects of the study design. All authors contributed to and critically reviewed the manuscript. DJB, CCC, PDK and SFWM made especially major contributions to the writing and editing.

## Additional information

Supplementary Information is available for this paper at [URL].

